## Supplemental figures and tables for "An Integrated Machine Learning Approach Delineates an Entropic Expansion Mechanism for the Binding of a Small Molecule to *α*-Synuclein"

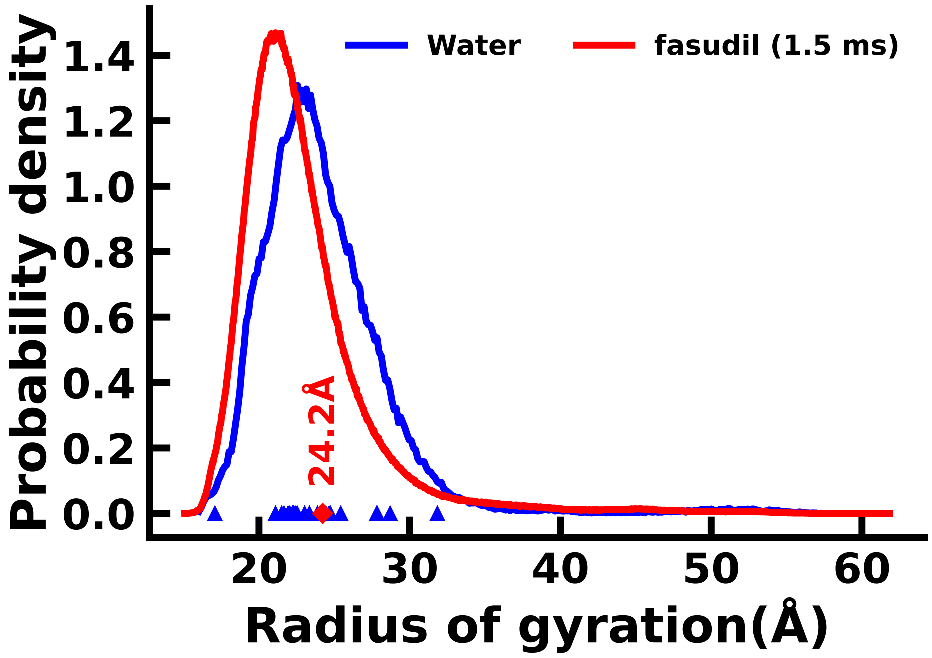


**Figure S1:** Distribution of the radius of gyration (Rg) of the $\alpha$S and $\alpha$S-Fasudil ensembles. The Rg values of the initial structures used for the $\alpha$S and $\alpha$S-Fasudil simulations are marked as symbols. The red diamond is the Rg of the starting structure of the $\alpha$S-Fasudil simulation and the blue triangle is the Rg of the starting structure of the $\alpha$S simulations

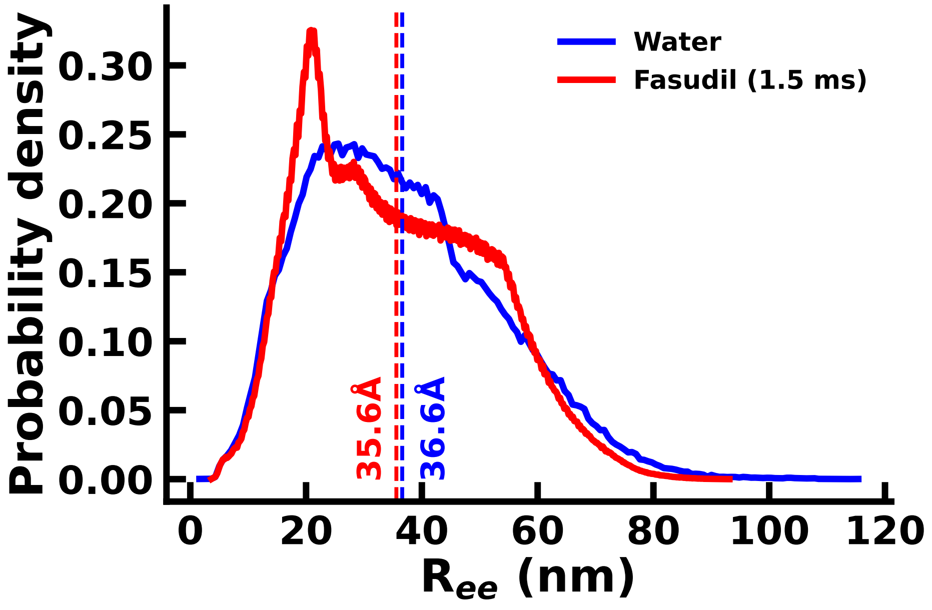


**Figure S2:** Distribution of end-to-end distance for the $\alpha$S and $\alpha$S-Fasudil ensembles. The mean values of the distributions are marked.

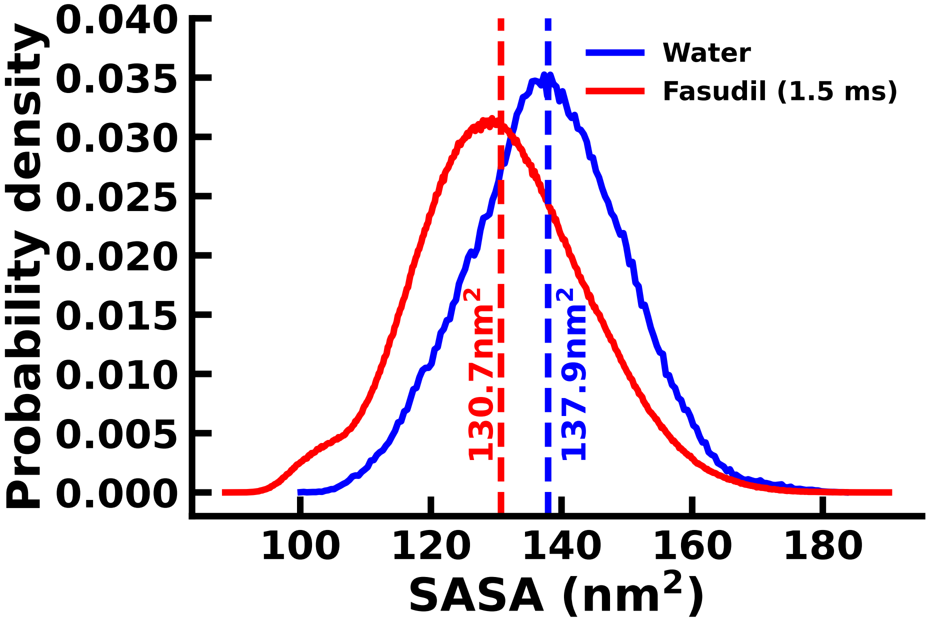


**Figure S3:** Distribution of SASA for $\alpha$S and $\alpha$S-Fasudil ensembles. The mean values of the distributions are marked.

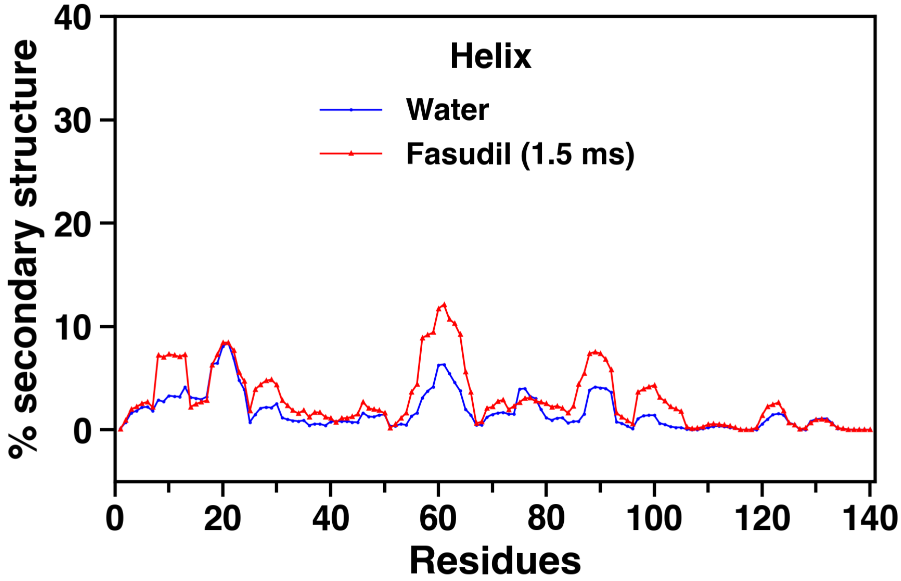


**Figure S4:** Residue wise percentage secondary structure of helix nature in the $\alpha$S and $\alpha$S-Fasudil ensembles.

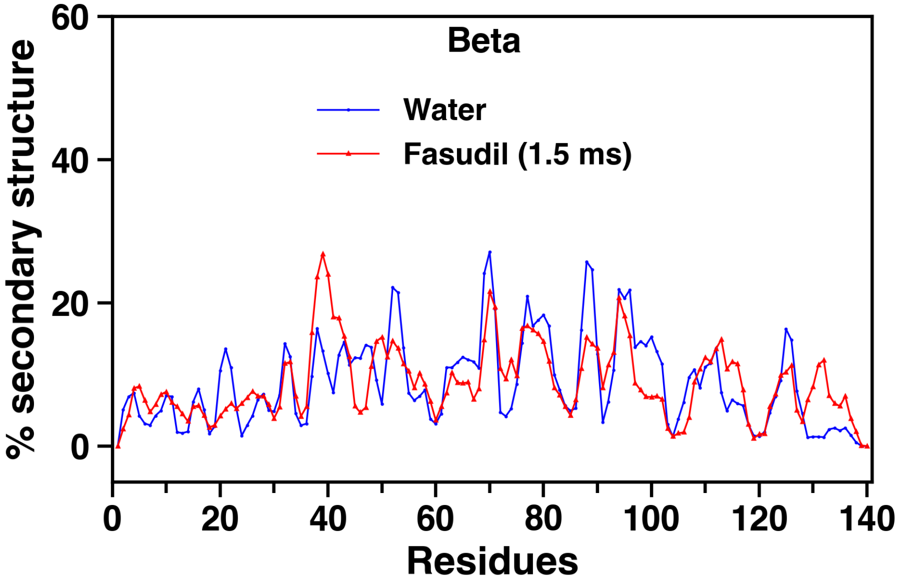


**Figure S5:** Residue wise percentage secondary structure of sheet nature in the $\alpha$S and $\alpha$S-Fasudil ensembles.

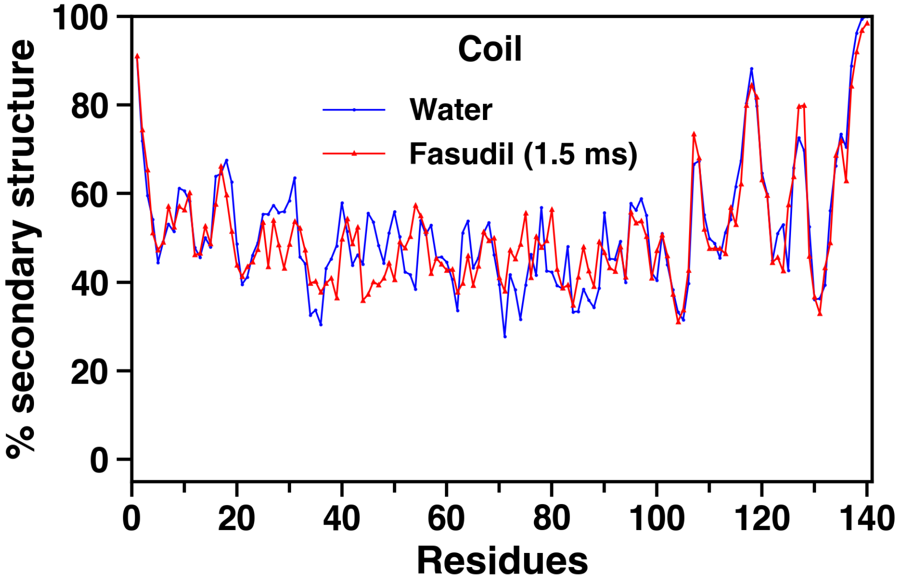


**Figure S6:** Residue wise percentage secondary structure of coil nature in the $\alpha$S and $\alpha$S-Fasudil ensembles.

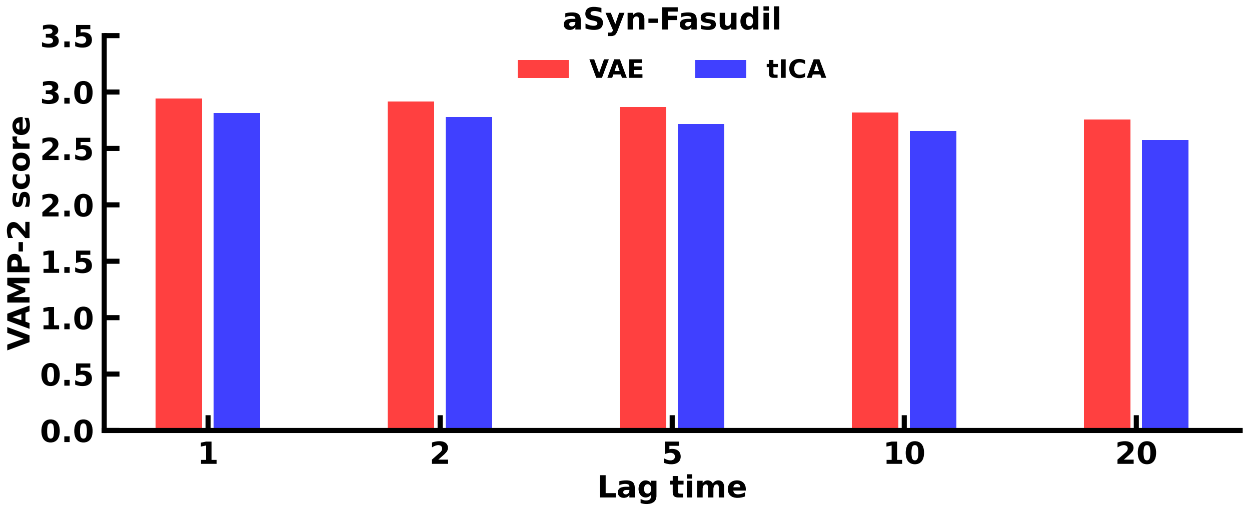


**Figure S7:** Comparison of VAMP-2 score from VAE and tICA of pairwise distances, reduced to two dimensions for the $\alpha$S-Fasudil simulation.

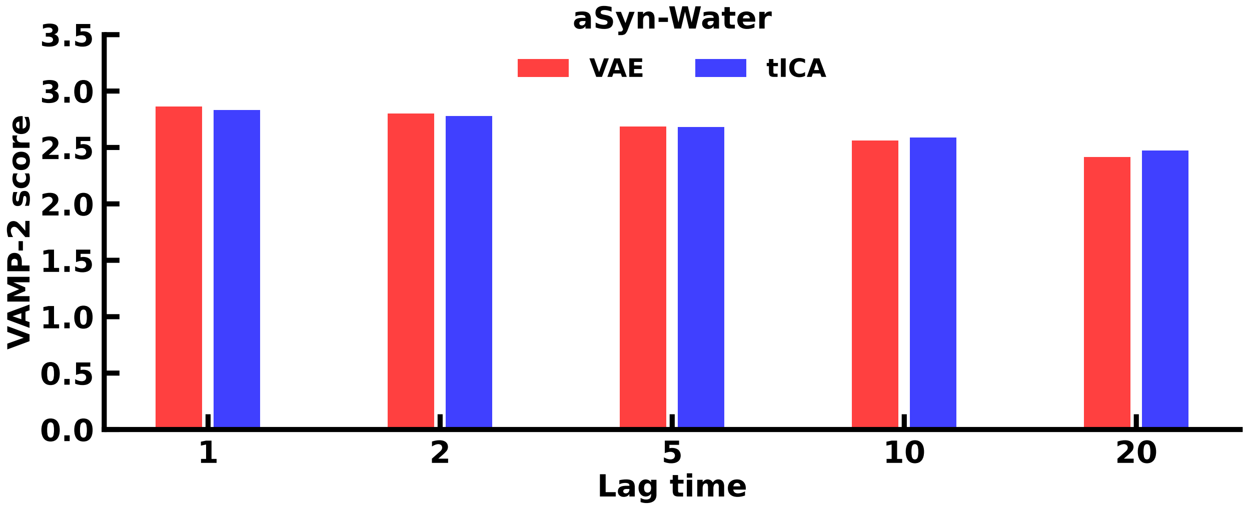


**Figure S8:** Comparison of VAMP-2 score from VAE and tICA of pairwise distances, reduced to two dimensions for the $\alpha$S simulation.

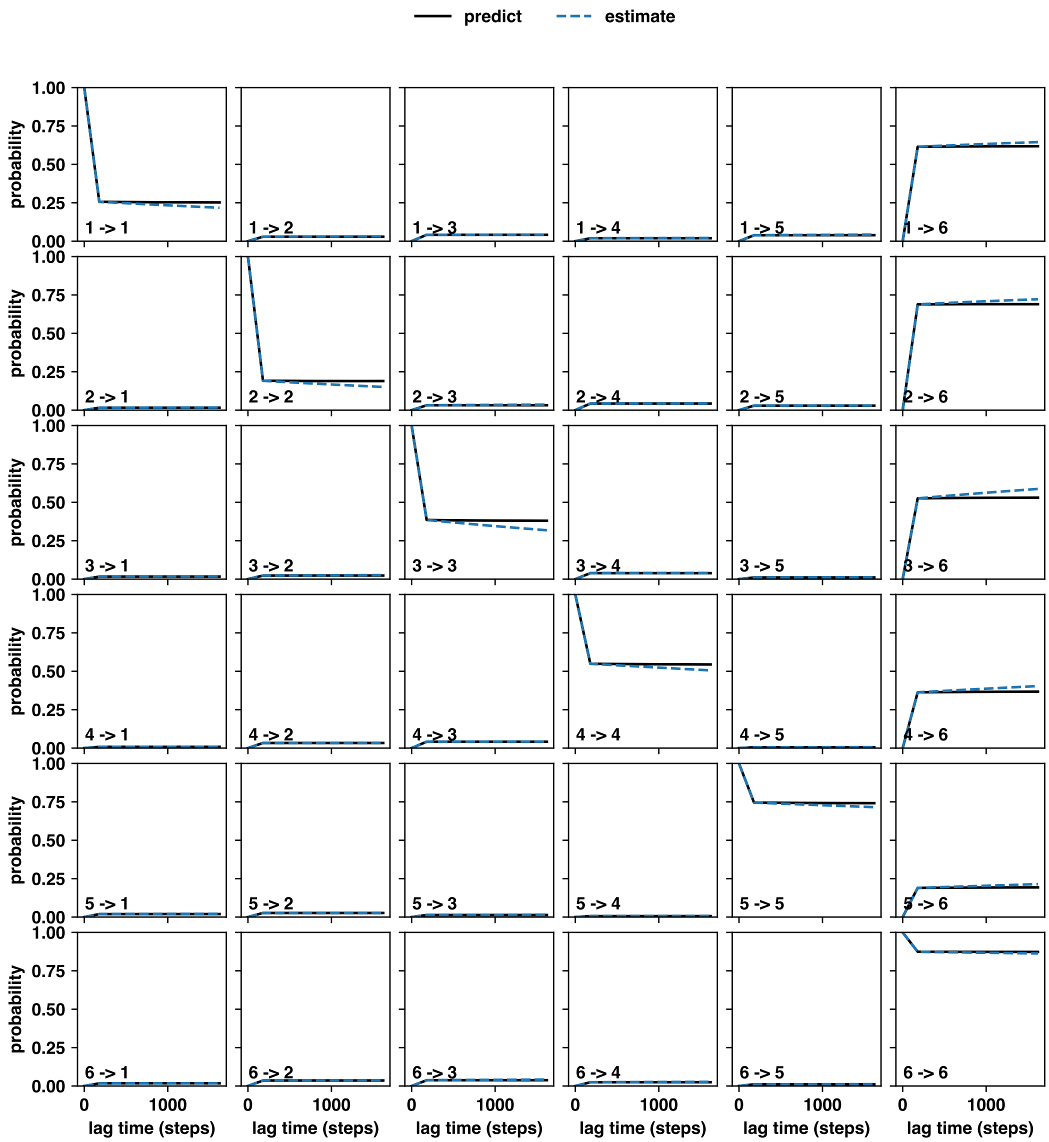


**Figure S9:** The Chapman-Kolmogorov test performed for the six state Markov State Model of the $\alpha$S-Fasudil ensemble

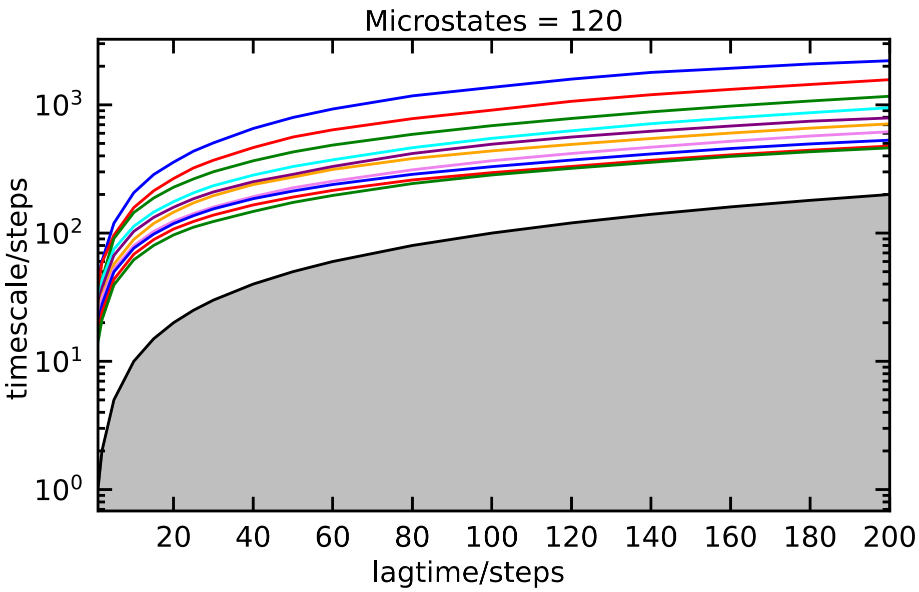


**Figure S10:** The implied timescale (ITS) plot $\alpha$S-Fasudil simulation for the block of 10 $\mu$s to 70 $\mu$s.

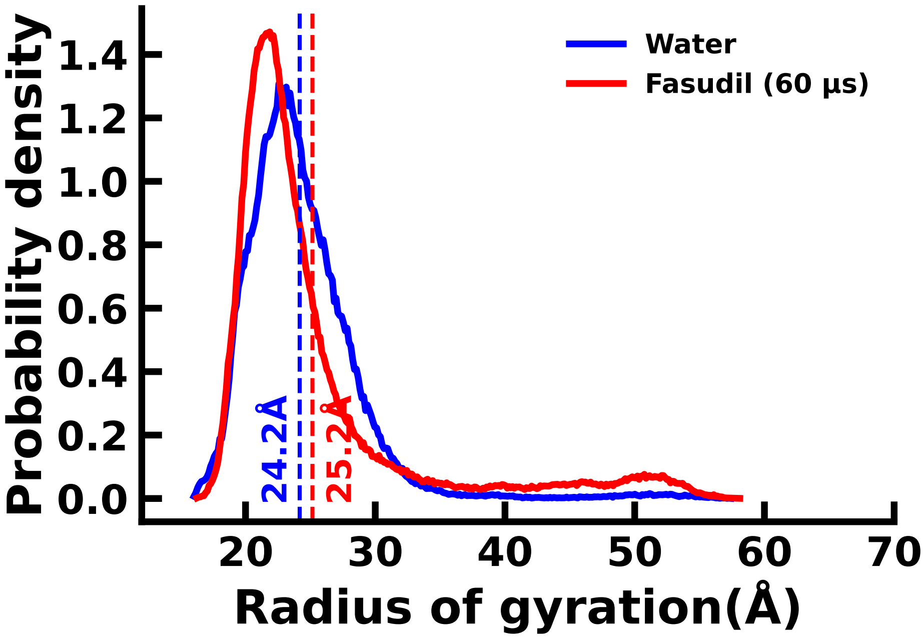


**Figure S11:** Distribution of Rg of the $\alpha$S ensemble and the 60 $\mu$s slice (10 $\mu$s-70 $\mu$s) of the $\alpha$S-Fasudil simulation trajectory.

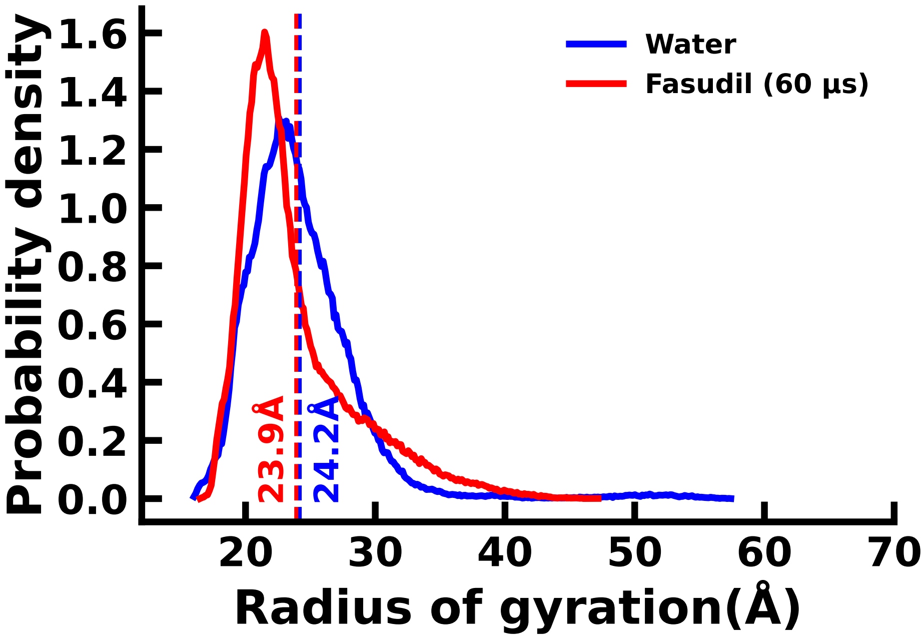


**Figure S12:** Distribution of Rg of the $\alpha$S ensemble and the 60 $\mu$s slice (966 $\mu$s to 1026 $\mu$s) of the $\alpha$S-Fasudil simulation trajectory.

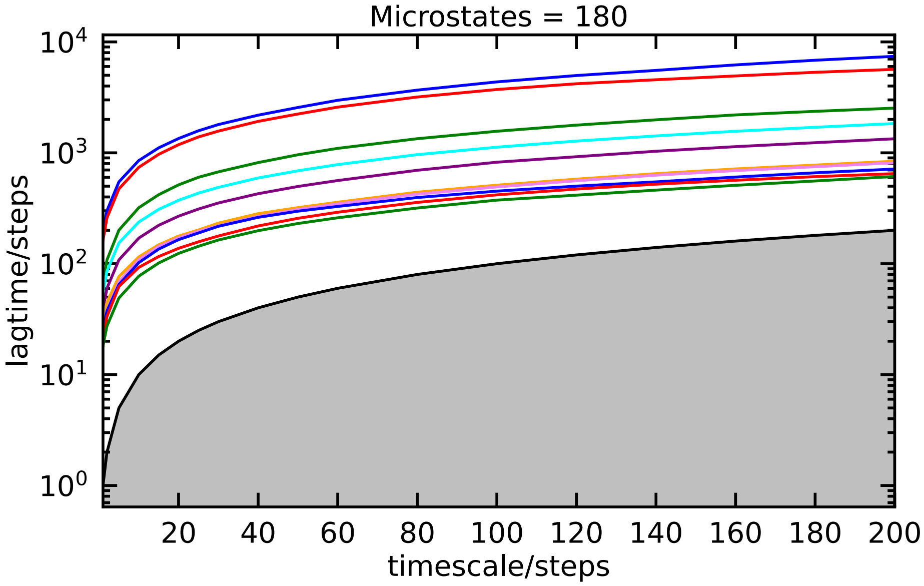


**Figure S13:** The implied timescale (ITS) plot $\alpha$S-Fasudil simulation for the block of 966 $\mu$s to 1026 $\mu$s. Here too we see 6 states.

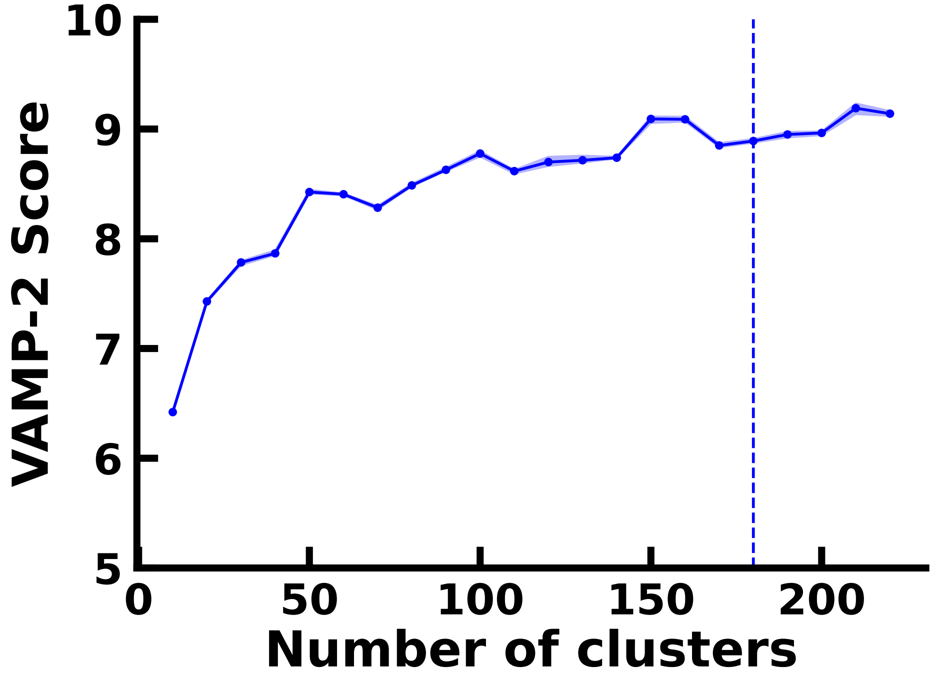


**Figure S14:** VAMP-2 score as a function of the number of microstates in for the $\alpha$S-Fasudi ensembles for a latent space of 4 dimensions.

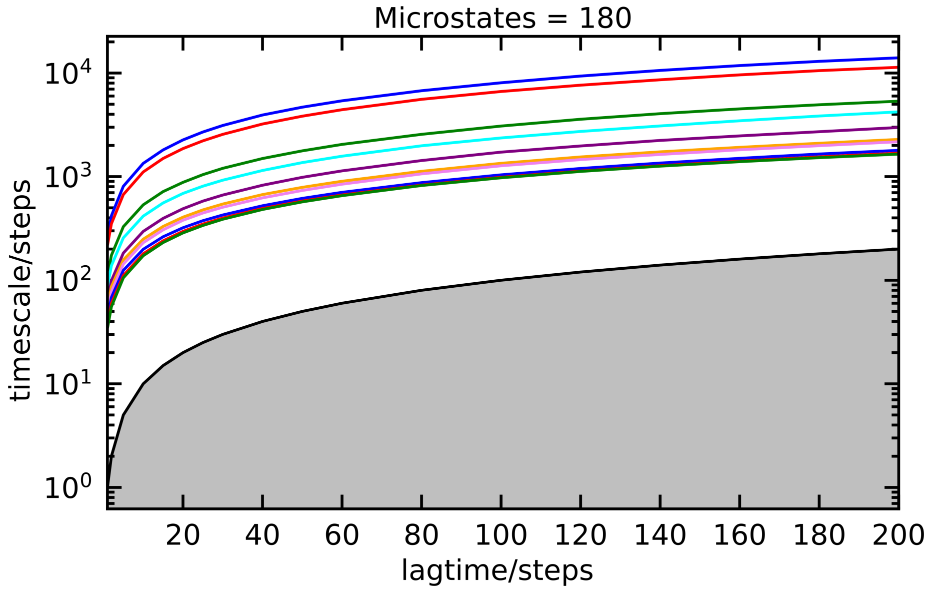


**Figure S15:** ITS plot of the $\alpha$S-Fasudi simulation using the 4 dimension of the VAE. Here too we see 6 states.

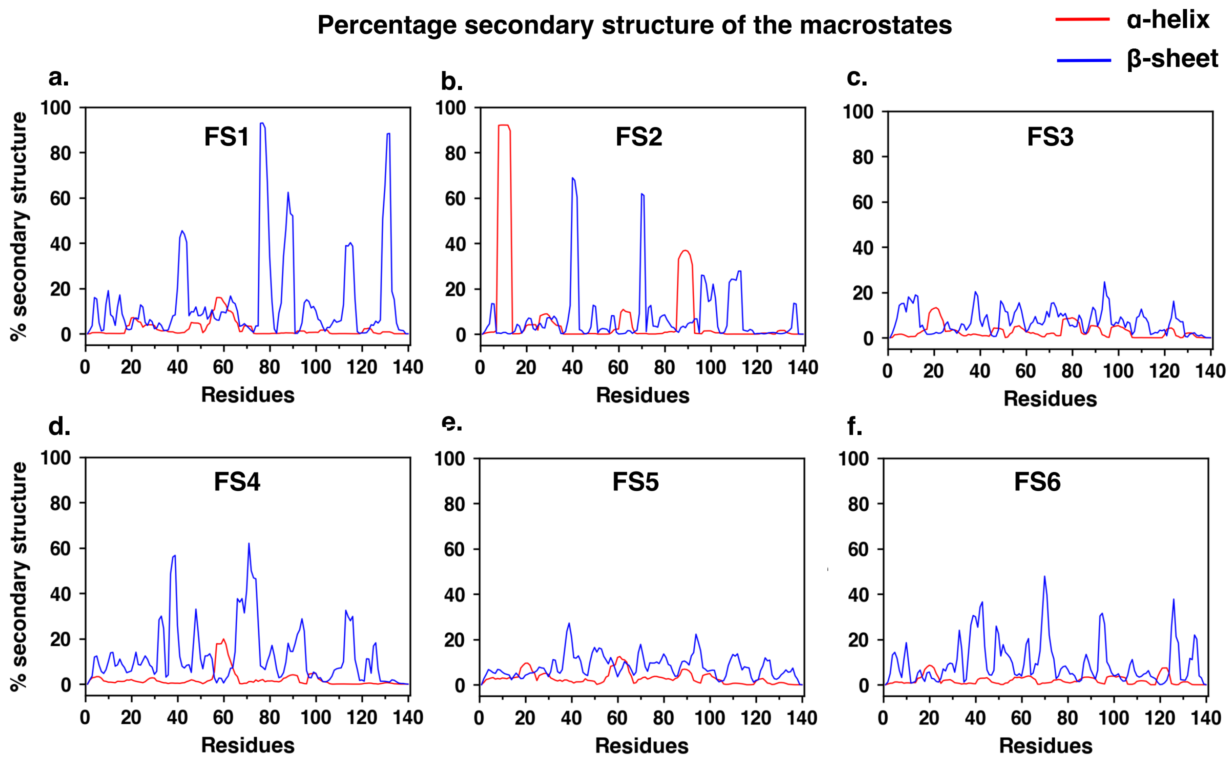


**Figure S16:** Residue-wise percentage secondary structure estimated for each of the six macrostates of $\alpha$S monomer simulated in the presence of fasudil.

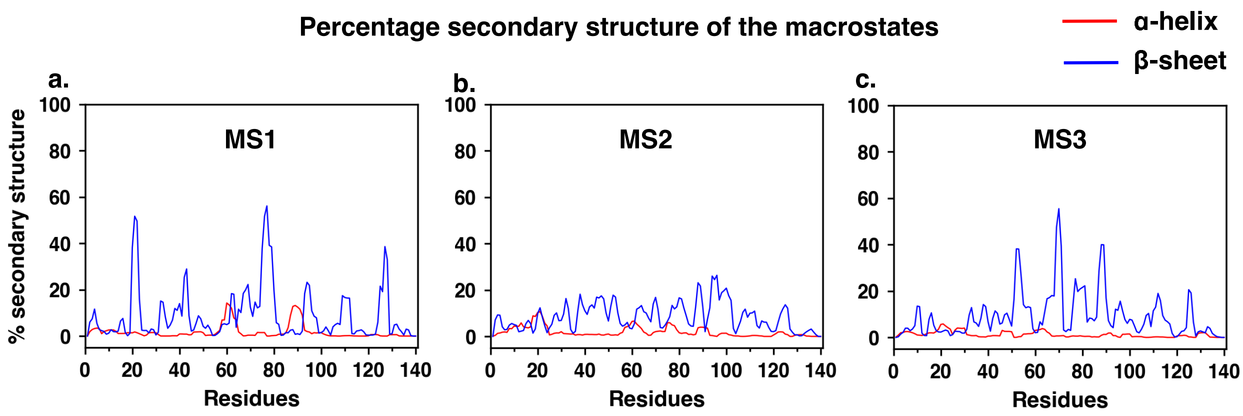


**Figure S17:** Residue-wise percentage secondary structure estimated for each of the three macrostates of $\alpha$S monomer simulated in neat water.

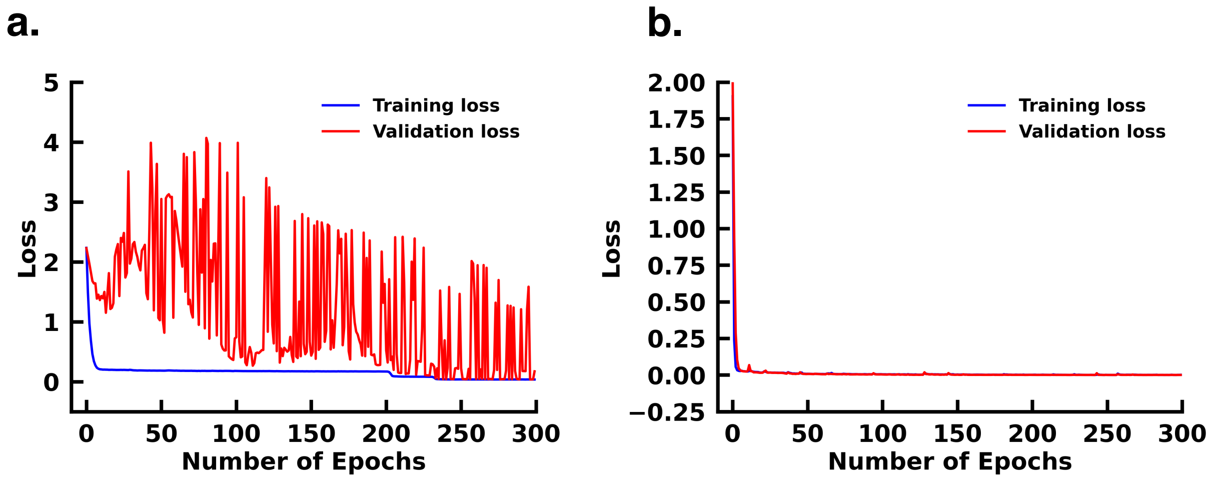


**Figure S18:** Loss profile of (a) CVAE indicating overfitting (b) DCVAE indicating no overfitting.

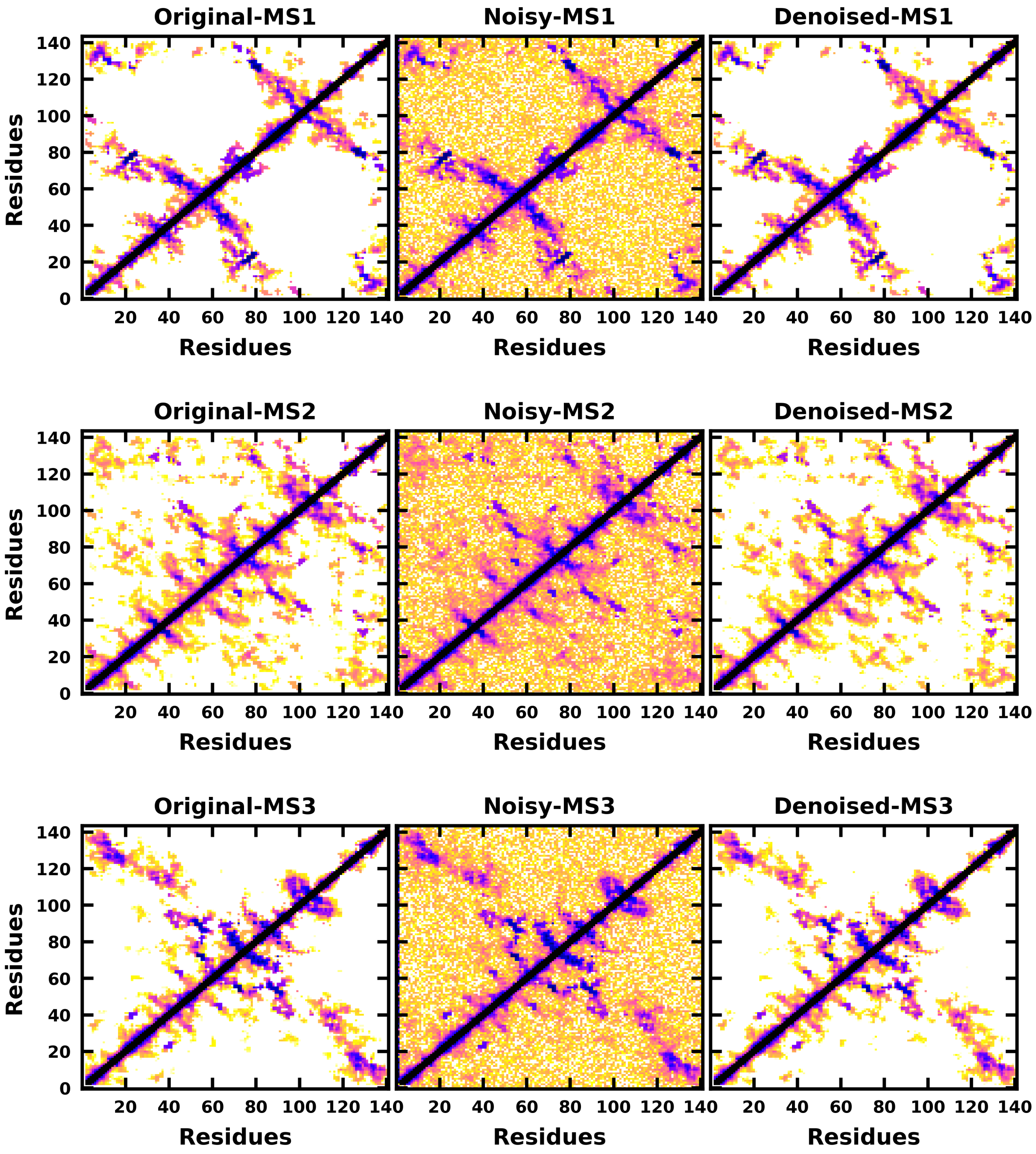


**Figure S19:** Denoised output of the contact map of the $\alpha$S macrostates.

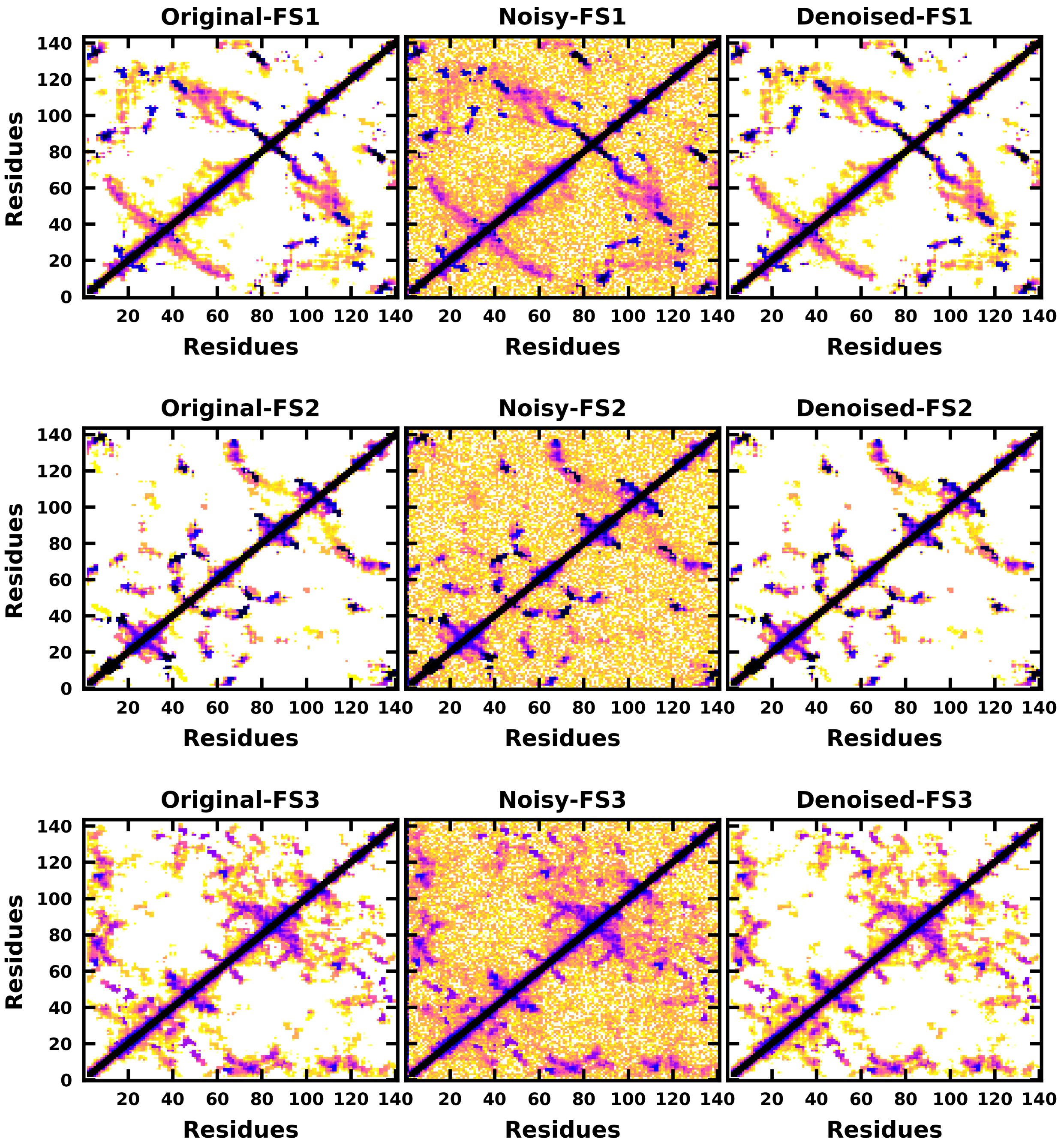


**Figure S20:** Denoised output of the contact map of the $\alpha$S-Fasudil macrostates FS1, FS2 and FS3.

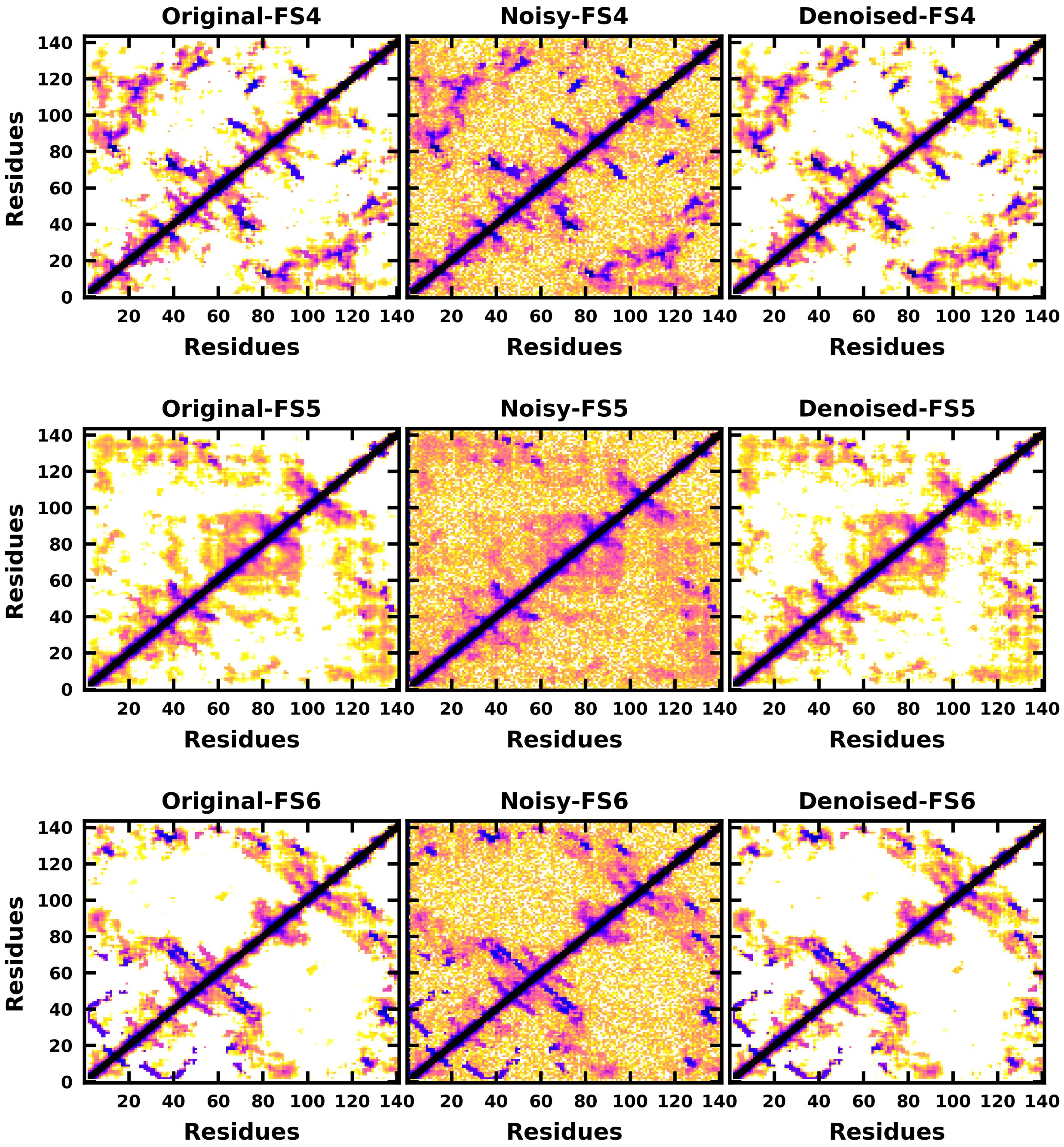


**Figure S21:** Denoised output of the contact map of the $\alpha$S-Fasudil macrostates FS4, FS5 and FS6.

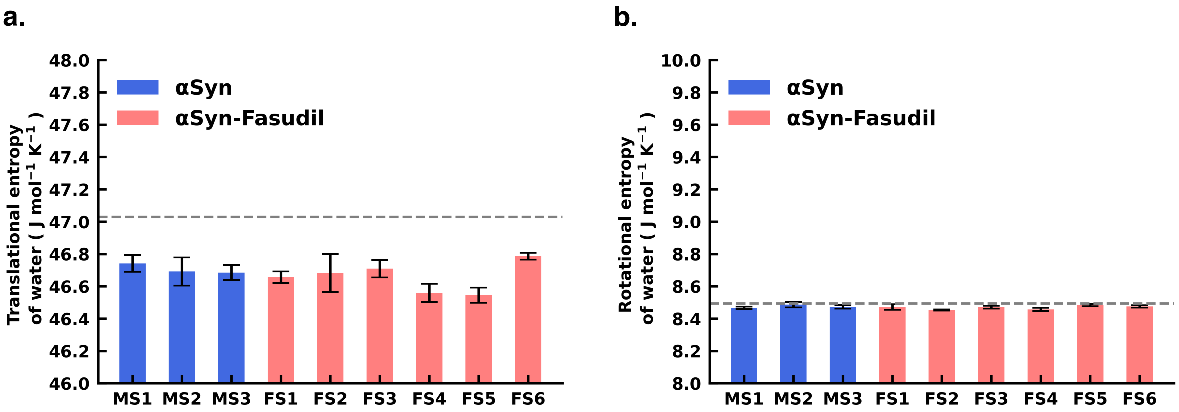


**Figure S22:** (a) Translational entropy of water and (b) Rotational entropy of water in $\alpha$S and $\alpha$S-Fasudil system.

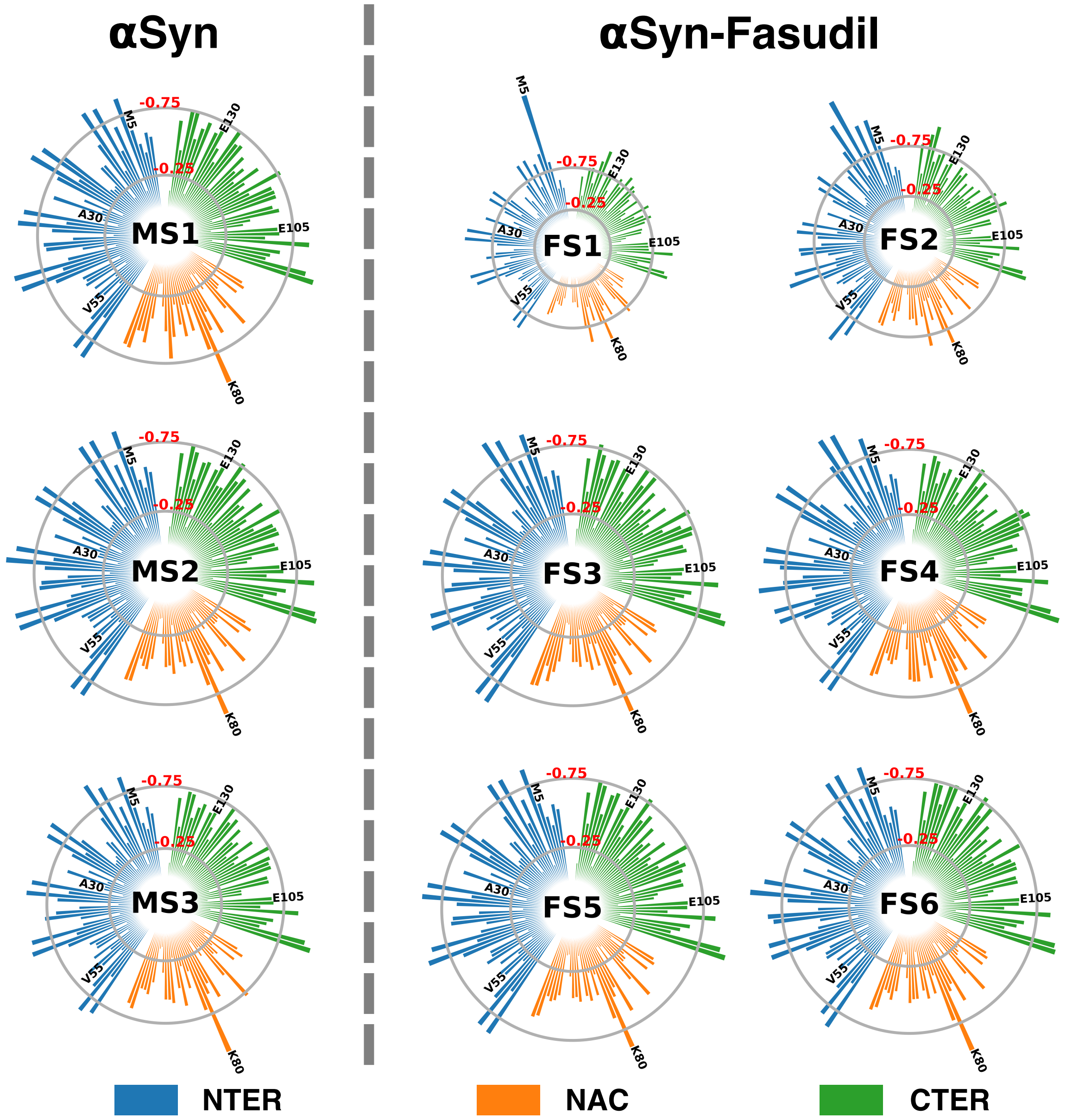


**Figure S23:** Residue-wise entropy of $\alpha$S states populated in water and in the presence of fasudil

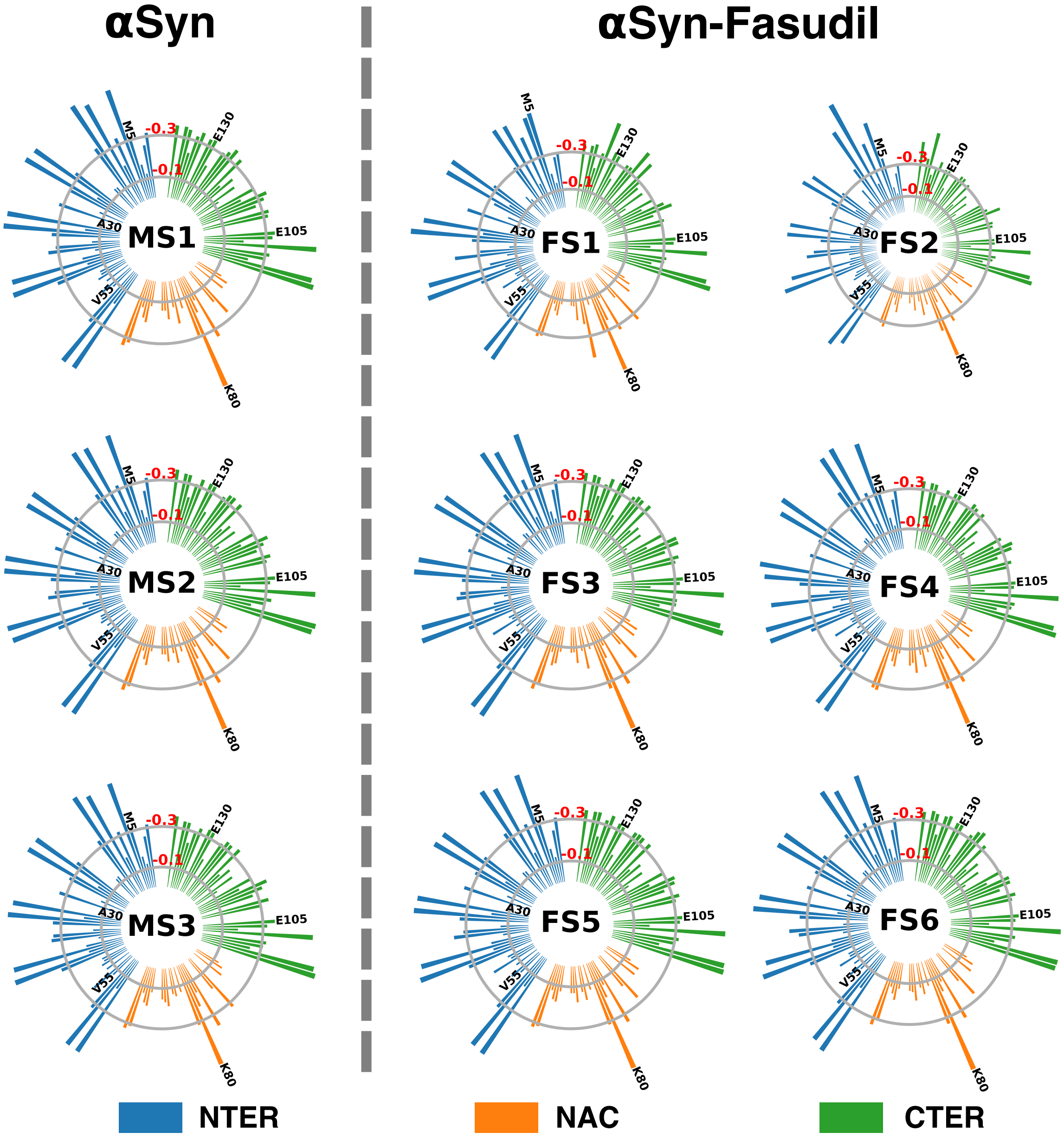


**Figure S24:** Residue-wise sidechain entropy of $\alpha$S states populated in water and in the presence of fasudil

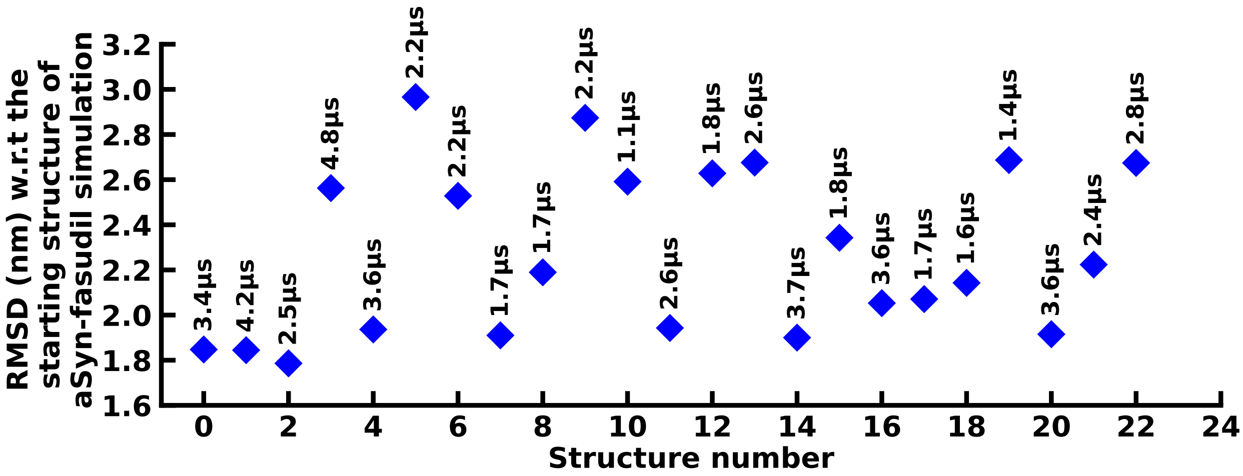


**Figure S25:** Each blue diamond label represents the RMSD (nm) w.r.t the starting structure of aSyn-fasudil simulation. The value associated with each diamond label corresponds to the timescale of simulation.

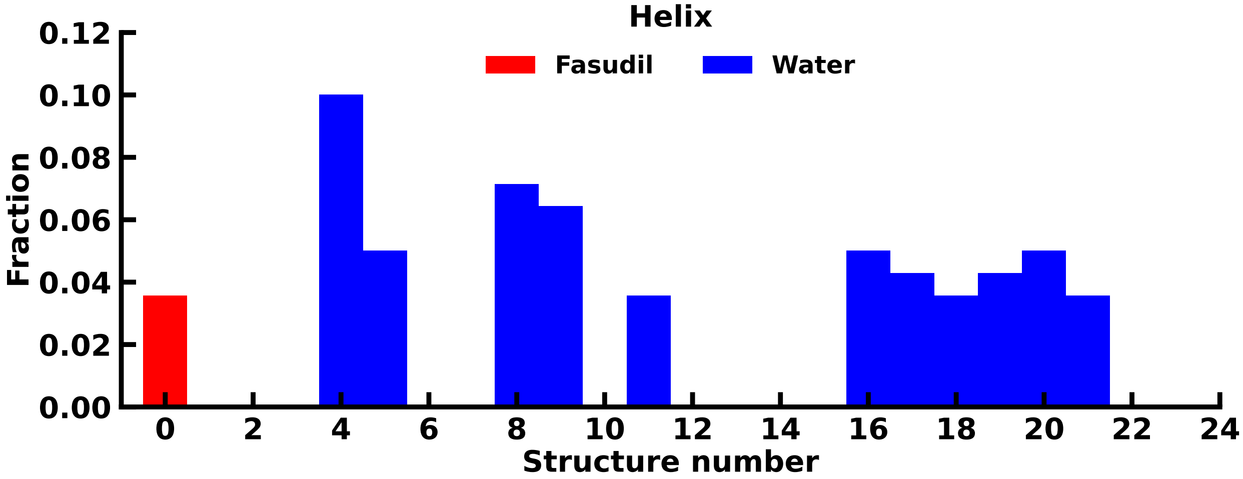


**Figure S26:** Fraction of helix content in the starting structures of the $\alpha$S-Fasudil and $\alpha$S simulations.

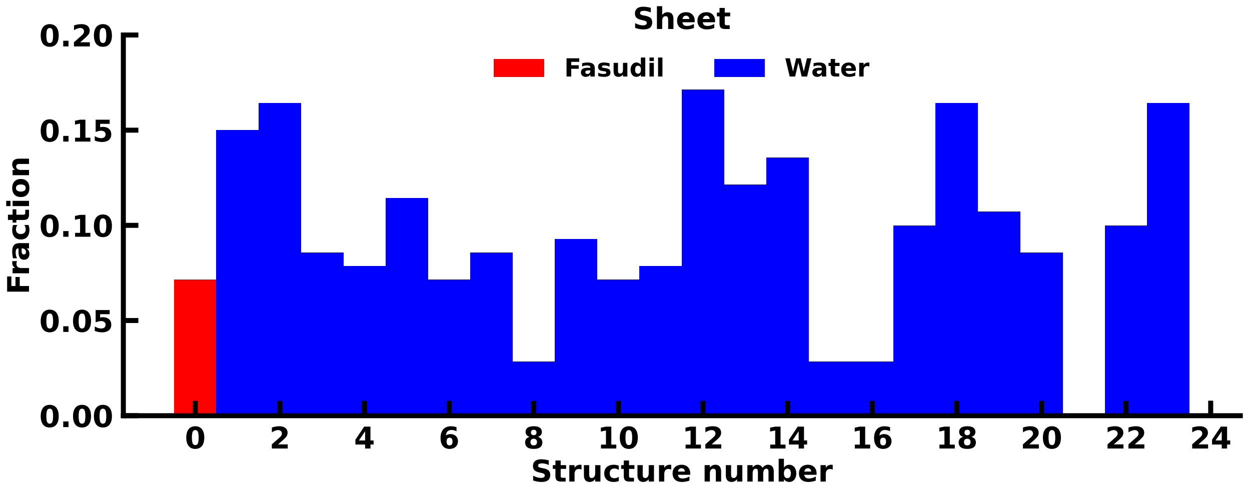


**Figure S27:** Fraction of $\beta$-sheet content in the starting structures of the $\alpha$S-Fasudil and $\alpha$S simulations.

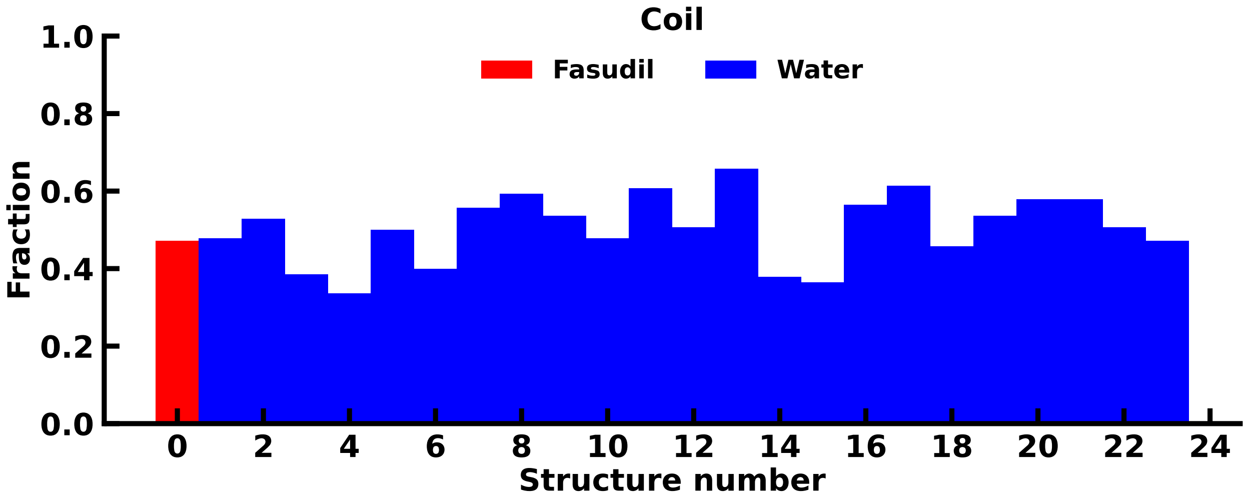


**Figure S28:** Fraction of coil content in the starting structures of the $\alpha$S-Fasudil and $\alpha$S simulations.

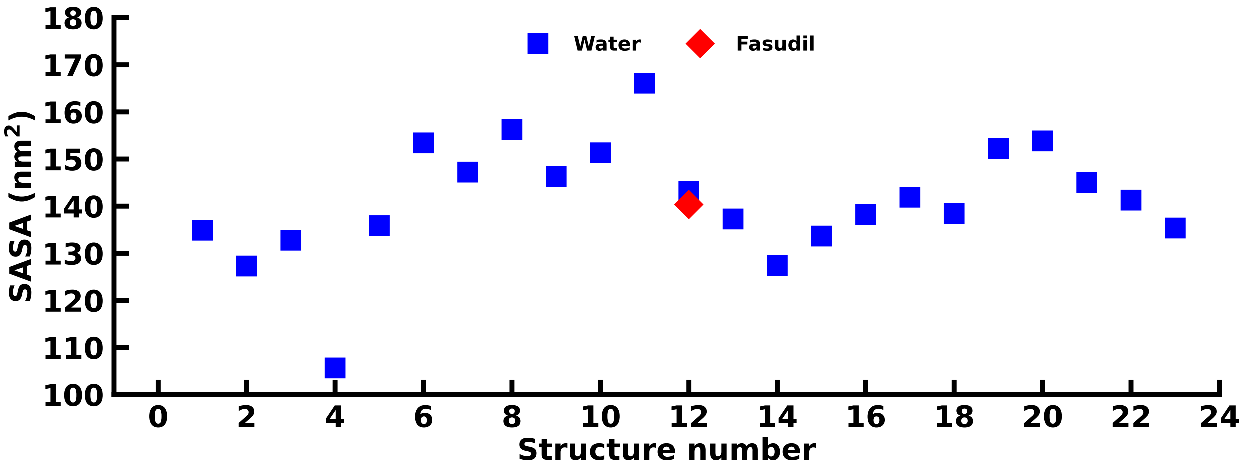


**Figure S29:** Total SASA of the starting structures of the $\alpha$S-Fasudil and $\alpha$S simulations.

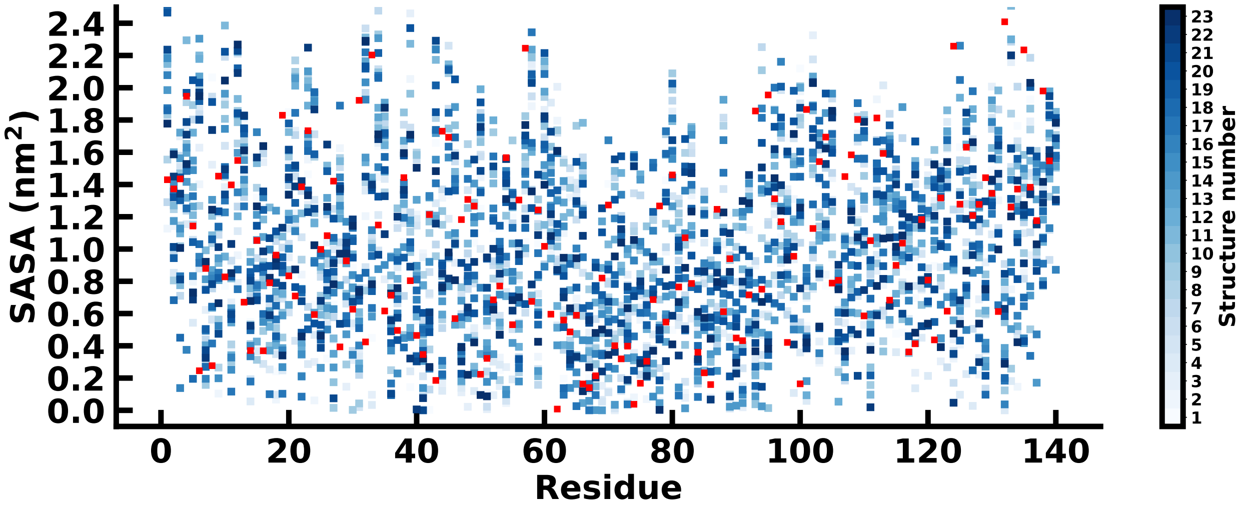


**Figure S30:** Residue wise SASA of the starting structure of the $\alpha$S-Fasudil (red square) and $\alpha$S (blue squares) simulations.

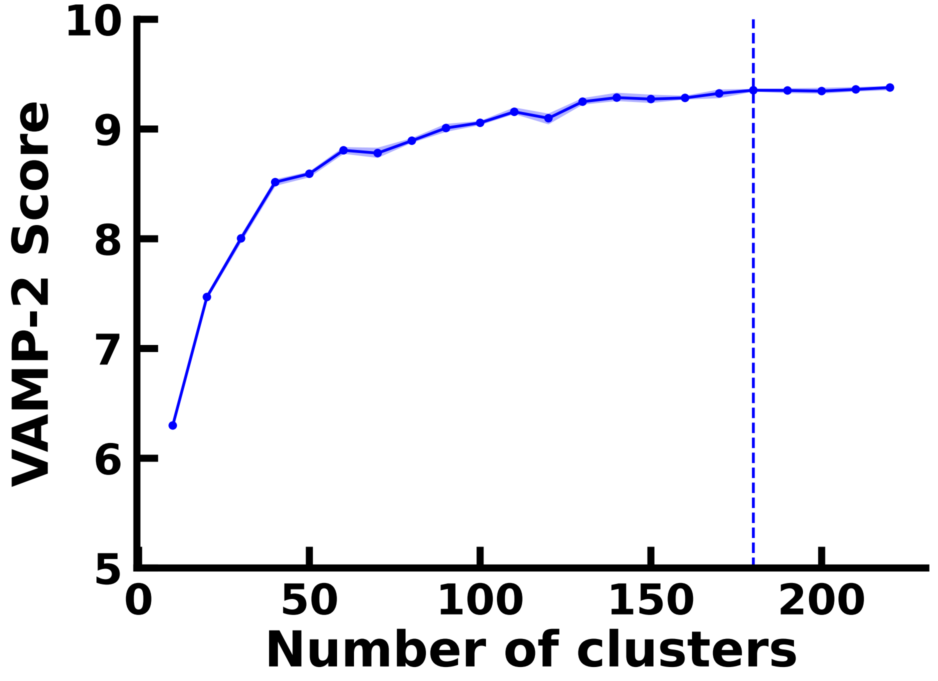


**Figure S31:** VAMP-2 score as a function of the number of microstates in for the $\alpha$S-Fasudil ensemble.

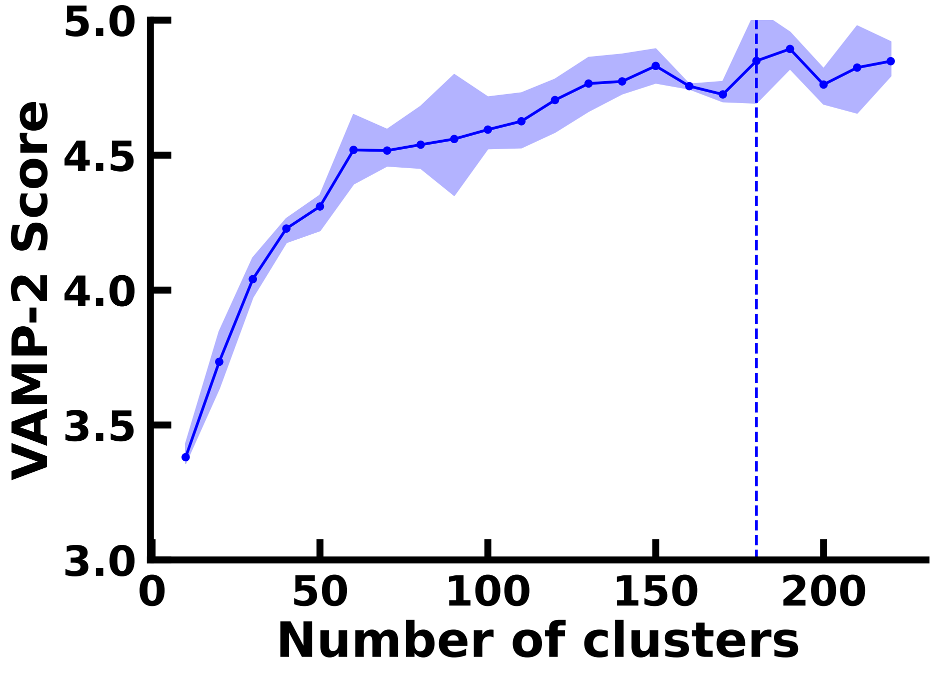


**Figure S32:** VAMP-2 score as a function of the number of microstates in for the $\alpha$S ensemble.

**Table S1:** The size of the simulation box for the 23 $\alpha$S simulations

| **Conformation number** | **Cubic Box size (nm)** |
|---|---|
| 1 | 12.14 |
| 2 | 11.33 |
| 3 | 13.55 |
| 4 | 10.41 |
| 5 | 10.97 |
| 6 | 10.44 |
| 7 | 10.10 |
| 8 | 11.44 |
| 9 | 10.57 |
| 10 | 10.75 |
| 11 | 11.34 |
| 12 | 10.35 |
| 13 | 13.38 |
| 14 | 10.08 |
| 15 | 10.38 |
| 16 | 11.78 |
| 17 | 11.92 |
| 18 | 9.37 |
| 19 | 10.15 |
| 20 | 11.16 |
| 21 | 11.24 |
| 22 | 11.46 |
| 23 | 12.00 |
